# Stability and critical transitions in mutualistic ecological systems

**DOI:** 10.1101/098046

**Authors:** Wenfeng Feng, Richard M. Bailey

## Abstract

Successful conservation of complex ecosystems, their function and associated services, requires deep understanding of their underlying dynamics and potential instabilities. While the study of ecological dynamics is a mature and diverse field, the lack of a general model that uses basic ecological parameters to predict system-level behaviour has allowed unresolved contradictions to persist. Here, we provide a general model of a mutualistic ecological community and show for the first time how the conditions for instability, the nature of ecological collapse, and potential early-warning signals, can be derived from the basic ecological parameters. We also resolve open questions concerning effects of interaction heterogeneity on both resilience and abundance, and discuss their potential trade-off in real systems. This framework provides a basis for rich investigations of ecological system dynamics, and can be generalised across many ecological contexts.

---

The relationships between various aspects of system complexity and system stability have recieved much attention in ecology[1, 2, 3], and in other fields (e.g. economic[4] and social[5] systems). Definitions of ecosystem stability are wide-ranging and, under deterministic dynamic models, extend across local asymptotic stability [6, 7], persistence[8, 9, 10], productivity (total abundance)[11], alternative steady states and catastrophic transitions[12, 13, 14, 15]; under stochastic dynamic models, these are extended further to include temporal community-level and species-level stability [16, 17, 18, 19, 20], and empirical signals of critical transitions [21, 22, 23, 24]. Similarly, the system properties associated with complexity and stability have a wide span, including biodiversity and connectance[25, 26], the strength and correlation of interactions[27, 28, 7], interaction asymmetry[29, 30, 31], and structural features such as degree heterogeneity and modularity[8, 32, 33, 28, 27, 11, 10, 34]. The type of ecological systems studied has progressed from random to competitive communities[16, 17, 35], to exploitative communities (food-webs)[36, 37, 10], mutualsitic communities[8, 32, 33, 28, 27, 11], and to competitive-exploitative-mutualistic mixed communities[38, 11]. Inconsistencies have emerged within this body of literature and an integrated framework that reveals how basic ecological parameters lead to these higher level phenomena is urgently needed [1, 39]. Here we develop such a framework using a general mutualistic model (two mutualistic groups of species, plants and pollinators for example, with competitive intra-group interactions). The ‘basic’ free parameters defining the model include per capita intrinsic growth rate *r* (which we adjust to simulate changes in environmental stress), per capita self-regulation strength *s*, competitive and mutualistic interaction strength (C, M), and resource handling time *h*. We focus initially on a mean field version of the full model[40], which preserves the essential features of a mutualistic system, but is simple enough to be mathematically tractable. We use this to identify necessary conditions for the presence of alternative stable states, then the transitions between them and their potential leading indicators.

## Presence and dynamics of alternative stable states

Understanding of alternative steady states, critical transitions and hysteresis within ecological contexts, in terms of the basic underlying ecological parameters, remains incomplete. We begin our analysis of these phenomena by first examining steady-state solutions, and the associated eigenvalue distributions of the relevant Jacobian matrices (definitions of ‘dot’ and ‘semicircle’ eigenvalues, λ*_d_*(J) and λ*_s_*(J) respectively, and the difference between them, the ‘spectral gap’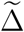, are provided in Methods and SOM). In cases where the dynamics are dominated by the ‘dot’ eigenvalue, the full set of basic free parameters can be conveniently reduced to a parameter space of three dimensions - handling time *h*, intrinsic growth rate *r*, and the ratio of total mutualistic strength to total competitive strength 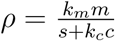 (see Methods). Solutions within this space can be partitioned into topologically equivalent regions (*strata*) of stability ([41]) (Fig.1), and these are strongly determined by handling time *h*. Only when *h* > 0 does bistability emerge (*H*_12_), with two basins of attraction representing positive abundance and zero abundance, separated by an unstable equilibrium (Fig.1). Indeed, this region (*H*_12_) provides a clear set of conditions for the appearance of alternative stable states in mutualistic ecosystems: (i) a non-linear functional response, reflected by the saturating coefficient for handling time, *h* > 0; (ii) mutualistic interactions that are stronger than competitive interactions, ρ > 1, and (iii) a negative intrinsic growth rate larger than a minimal value 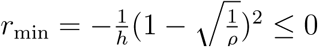

**Figure 1.**
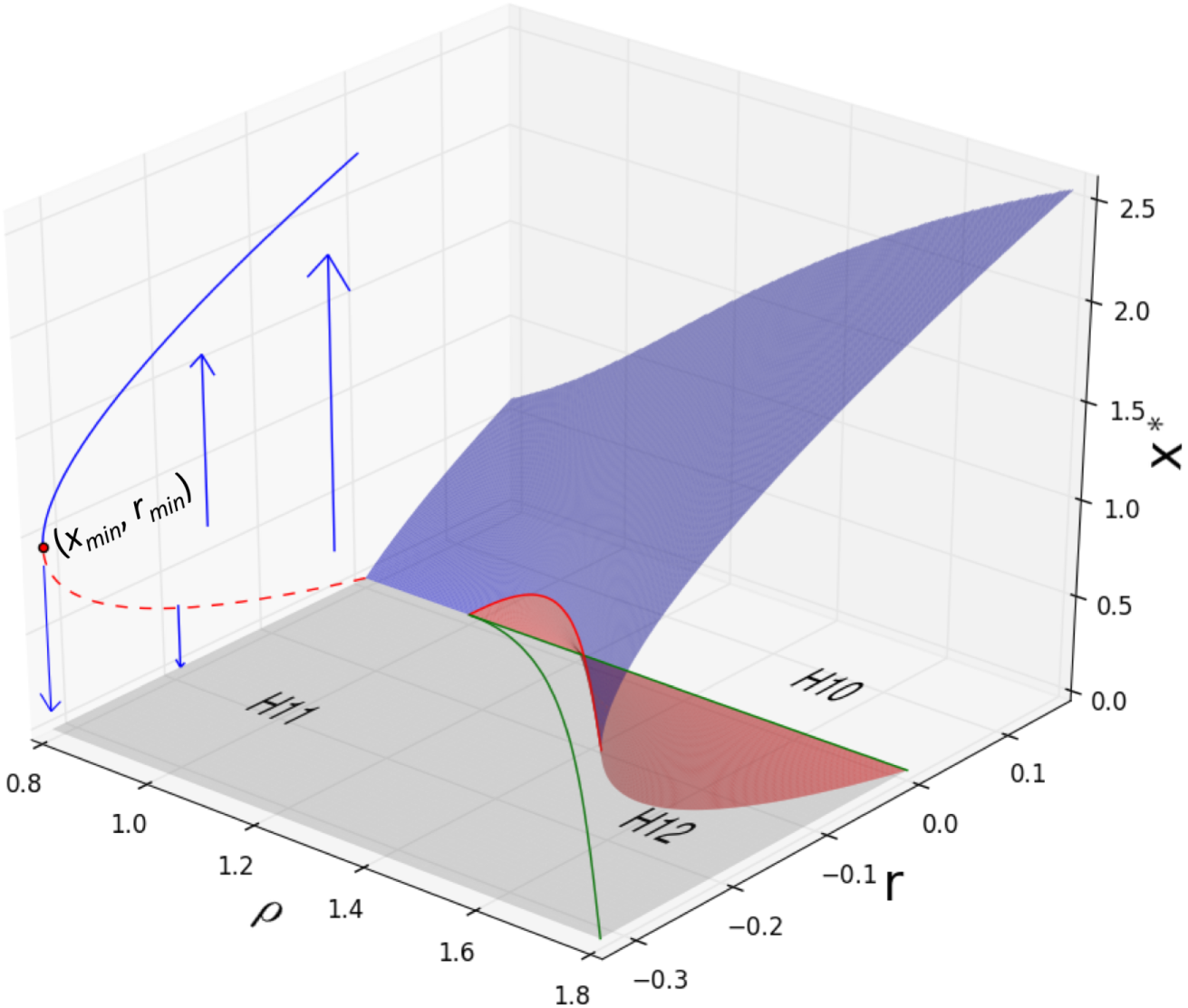
Parametric portrait and phase portrait from the mean filed model. For the case where ‘dot’ eigenvalues dominate, the parametric plane of intrinsic growth rate *r* and relative mutualistic strength *ρ*, is partitioned into three strata: *H*_10_ - globally stable at a positive equilibrium, *H*_11_ - globally stable at zero, and *H*_12_ - bistable both at a positive equilibrium and at zero. The phase portrait of *H*_12_ stratum is a cusp diagram with three sheets of equilibrium abundances **x*\** (blue, red and grey respectively), a fold curve (red solid line) and a cusp-shaped curve (green lines) which form the border of *H*_12_. A slice through the phase portrait of the *H*_12_ stratum, where *ρ* = 1.8, is shown on the r and *x*\* plane, in which stable states (blue line) intersect with unstable states (dashed red line) at the specific (*r_min_*, *x_min_*) point where critical transitions happen. For all results shown, *h* = 0.2.

Dynamics within stratum *H_12_* are clearly of particular interest and this space can be further described by the cusp structure [42] (Fig.1) using two control parameters *r*, *ρ* and one state variable - the equilibrium species abundance *x*\*. All values of *x*\* shown are equilibrium states and these comprise three sheets: an upper sheet of locally stable positive equilibrium abundances (*x*\* > 0); a lower sheet of locally stable zero abundance (*x*\* = 0); and (iii) the middle sheet of unstable equilibrium abundance (the border between the two ‘basins of attraction’). Critical transitions can happen where the upper and middle sheets intersect (the‘fold curves’ shown as red solid lines in Fig.1). When the two fold curves are projected back onto the plane of the control surface, the result is a cusp-shaped curve (green line in Fig.1) which forms the border of parameter stratum *H*_12_. If we suppose the system is forced through the parameter strata by varying the growth rate, *r*, the resultant dynamics (including the possibility of critical transitions) depends not only on *r* but on the previous state (the sheet occupied). Consequently, hysteresis can be observed in these results, which is measured here by the width of parameter stratum *H*_12_ (the absolute value of *r*_min_) and relevant partial derivatives of *r*_min_ show the width of *H*_12_ to be negatively proportional to the inverse square of handling time *h*, and positively related to the relative mutualistic strength *ρ* (SOM). Hence, the stronger the mutualistic interactions and the greater the efficiency of resource handling, the wider the parameter stratum *H*_12_, and the greater the extent of hysteresis. A consequence of this is that more strongly mutualistic systems tolerate what can be interpreted as harsher conditions (lower *r*), but this capability is associated with a ‘cost’ of a greater difficulty in recovering from collapse. While the occurence of such critical transitions has been studied in many types of systems (e.g. [39, 5, 4]), the precise nature of these transitions has recieved less attention, and this is the focus of the following section.

## Nature of critical transitions

We find that critical transitions in mutualistic systems follow one of two possible forms, and which of these occurs is strongly determined by whether the ‘dot’ or ‘semicircle’ eigenvalue dominates (reaches zero first) as the transition is approached; a similar claim has recently been made for physical many-body systems[43]. Where the ‘dot’ eigenvalue dominates, the associated leading eigenvector is an identity vector 1. The effect of reduced *r* (increased environmental stress) is therefore the same for each species, resulting in similar trajectories, and simultaneous population collapse during a critical transition. We name this a *consistent* transition (see Fig.2a,b). When the ‘semicircle’ eigenvalue dominates, the leading eigenvector has mixed negative and positive values. At the point of transition (which occurs at larger values of *r* compared to *consistent* transitions, equivalent to less harsh environmental conditions), the impacts on individual species are different in magnitude and even in sign, causing a range of trajectories. Some species abundances increase while others decrease or even collapse. We name this a *splitting* transition (see Fig.2a,b), and note that these have also been observed in numerical experiments [33]. The occurrence of *splitting* transitions (compared to *consistent* transitions) increases with the difference between the ‘semicircle’ eigenvalue and the ‘dot’ eigenvalue (—Δ^˜^), and this can be understood in terms of a new parameter α, the ratio between the ratio of the mutualistic spectral gap to competitive spectral gap Δ and the ratio of mutualistic strength to competitive strength *ρ*, i.e. 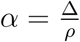. As *α* combines several parameters, we can evaluate their effect: the probability of a splitting transition increases with competitive strength *c*, decreases with both self-regulation (*s*) and the number of mutualistic interactions *k_m_*, and is unaffected by mutualistic strength *m* (SOM).

**Figure 2.**
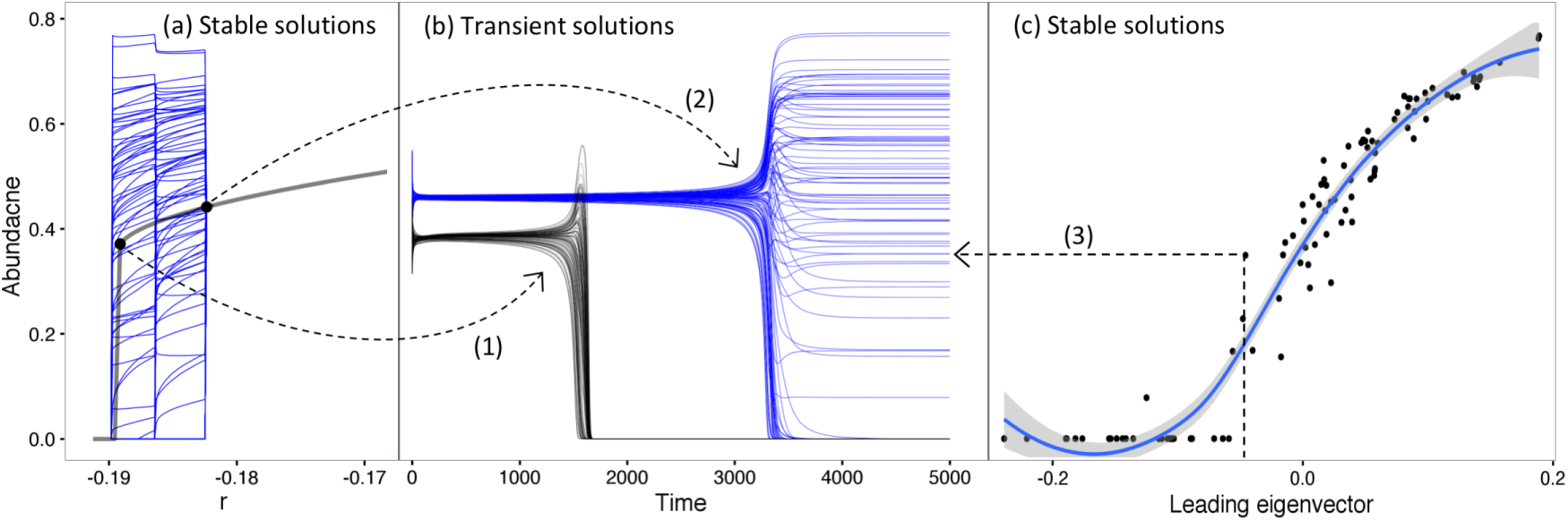
Progression from consistent transition to splitting transition under deterministic dynamics. The aim here is to isolate and demonstrate the effect of a on the nature of critical transitions. Starting with parameters *n* = 100, *k_m_* = 5, *k_c_* = *n*/2 — 1, *m* — 0.5,*c* = 0.004, *s* = 1, *h* = 0.5 (giving *ρ* ≈ 2.09, *α* ≈ 1.4), *ρ* is kept fixed and *α* is decreased by increasing c and decreasing *s* concurrently. Equilibrium abundance *x*\* is then not directly driven by changes in *α* but the nature of the critical transitions is affected. For *α* > 1, consistent transitions occur, and as *a* decreases below 1 the probability of splitting transition increases: line (1) connects the stable solution in (a) to the transient route to the stable solution in (b) for *α* ≈ 0.94); line (2) is for *α* ≈ 0.7), where splitting transitions are more probable. Splitting of abundances is determined by the corresponding elements of the leading eigenvector of the Jacobian matrix at the critical point, and (c) shows this relationship, exemplified by line (3). A polynomial fit is included only as a visual aid.

## Early warning of critical transitions

Of practical relevance is the question of empirically assessing proximity to critical thresholds, and here, whether this is different for the two identified types of transitions. We extend our analysis to transient stochastic dynamics and focus on two empirically observable aspects of species abundance time-series: variance of the total abundance of *n* species, *V^c^*, and the degree of synchronization among species η, which measures correlation through time of all viable species abundances (see SOM for details and Fig.2); both of these measures have been used previoulsy as ‘early-warning’ indicators ([21, 22]). We find that *V^c^* is negatively and inversely related to the ‘dot’ eignevalue λ*_d_*(J) and that *V^c^* therefore is indeed expected to increase sharply towards consistent-type critical transitions (as λ*_d_*(J) approaches zero). Moreover, because the partial derivative of λ*_d_*(J) also increases (as *r* decreases), the derivative of *V^c^* also increases, providing further indication of threshold proximity. At the same time, the spectral gap also increases, leading to significantly greater synchronization (η) among species (see SOM for details, and Fig.2). However, for *splitting* transitions, the ‘dot’ eigenvalue remains negative and changes relatively slowly with *r* (as it is relatively far from zero). Consequently, total variance *V^c^* is low compared with *consistent* transitions and the derivative of *V^c^* has no significant increasing trend, since it only increases markedly when λ*_d_*(J) is close to zero. Similarly, increases in synchronisation (η) are very weak prior to *splitting* transitions. As the *splitting* transition occurs, species abundances become un-correlated (Fig.2 and SOM). Thus, unlike the lead-up to *consistent* transitions, prior to *splitting* transitions the total variance *V^c^*, its derivative, and the synchronization among species, are not expected to provide clear empirical signals that anticipate critical transitions. Changes in the relevant eigenvalues have a marked effect on stability, and on the nature and predictability of critical transitions. We turn now to the influence of network structure on these, and on another closely-linked ecological property, total abundance.

## Effects of degree heterogeneity

The importance of structural aspects of ecological systems in determining higher level properties, such as stability, is well-recognised [8, 32, 33, 28, 27, 11, 10, 34]. For mutualistic systems, the degree of heterogeneity in the number of inter-species links is strongly correlated to nestedness [44], and defines the frequency distribution of more or less specialist/generalist species. Conflicting evidence exists however for the effect of this heterogeneity on stability (specifically *persistence,* the survival of all species, and *resilience,* the post-perturbation recovery rate) and also on total abundance [9, 10, 8, 11]. To study heterogeneity effects, we release the restriction of all species having the same number of mutualistic interactions (*k_m_*) and specify a vector describing the number of mutualistic interactions for each species (k_*m*_), within which the total mutualistic strength is held constant (SOM). We approximate the structure of real mutualistic systems by creating interaction networks with power-law (‘scale-free’) degree distributions (with exponent γ, noting the relationship to the ‘fitness coefficient’, *β*, by γ = 1/*β* +1 [45]; SOM). As a starting point we assess how the abundance of individual species with different numbers of interactions *(k_m_)* is affected by the heterogeneity of their interaction network (*β*), finding that species with more connections (higher *k_m_*) have greater abundance, and that this is increased further as the heterogeneity of their network is increased (larger β; Fig.4a). As might be expected for the community as a whole, mean abundance is greater when growth rates (*r*) are higher and mutualism is stronger (larger *ρ*). However, the relationship of mean abundance to heterogeneity (β) under changing *r* and *ρ* is more complex: a critical *r* value exists (which decreases as *ρ* increases), above/below which heterogeneity increases/decreases mean abundance (Fig. 4b). The effect of heterogeneity on total abundance therefore depends on both the strength of mutualism (*ρ*) and on the intrinsic growth rate (*r*), as shown in Fig.4a-c. Previous results that heterogeneity increases total abundance [11] are therefore confirmed, but seen to be special cases of weak mutualism and intrinsic growth rates below the identified critical value.

**Figure 3.**
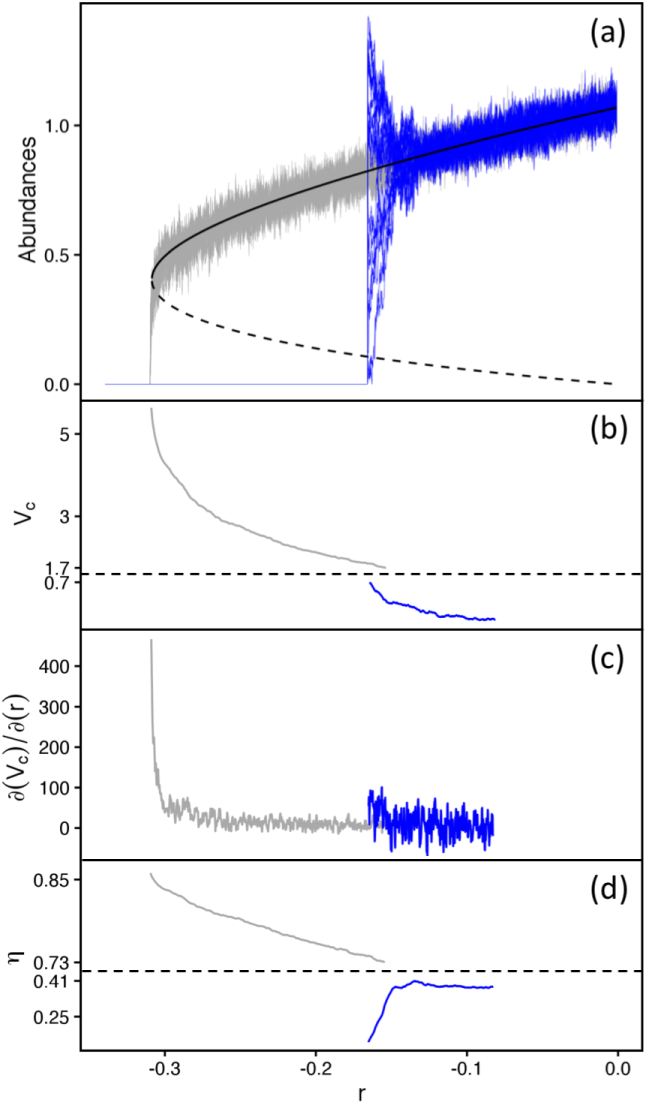
Early warnings of a consistent and a splitting transition. Transient stochastic simulations showing total species abundances under progressively reduced *r* and associated ‘early-warning’ signals for *consistent* (grey) and *splitting* (blue) transitions: (a) abundance of each of *n* species; (b) total variance, *V^c^* (horizontal dashed line in (b) and (d) indicates a break in the vertical axis); (c) the partial derivative of *V^c^* ∂(*V^c^*)/∂(*r*); and (d) the synchronization among species *η* (see SOM for full definitions). The values of *ρ* were kept fixed for both consistent and splitting cases, yielding the same mean equilibrium abundances before the transitions, in order to exclude other factors. Simulation conditions: *n* = 20, *k_m_* = 4, *k_c_* = *n*/2 — 1, *m* = 0.8, *h* = 0.5 and standard deviation of stochastic noise *σ* = 0.04 for all simulations, *c* = 0.02, *s* = 1 for consistent transitions, *c* = 0.08, *s* = 0.46 for splitting transitions.

**Figure 4.**
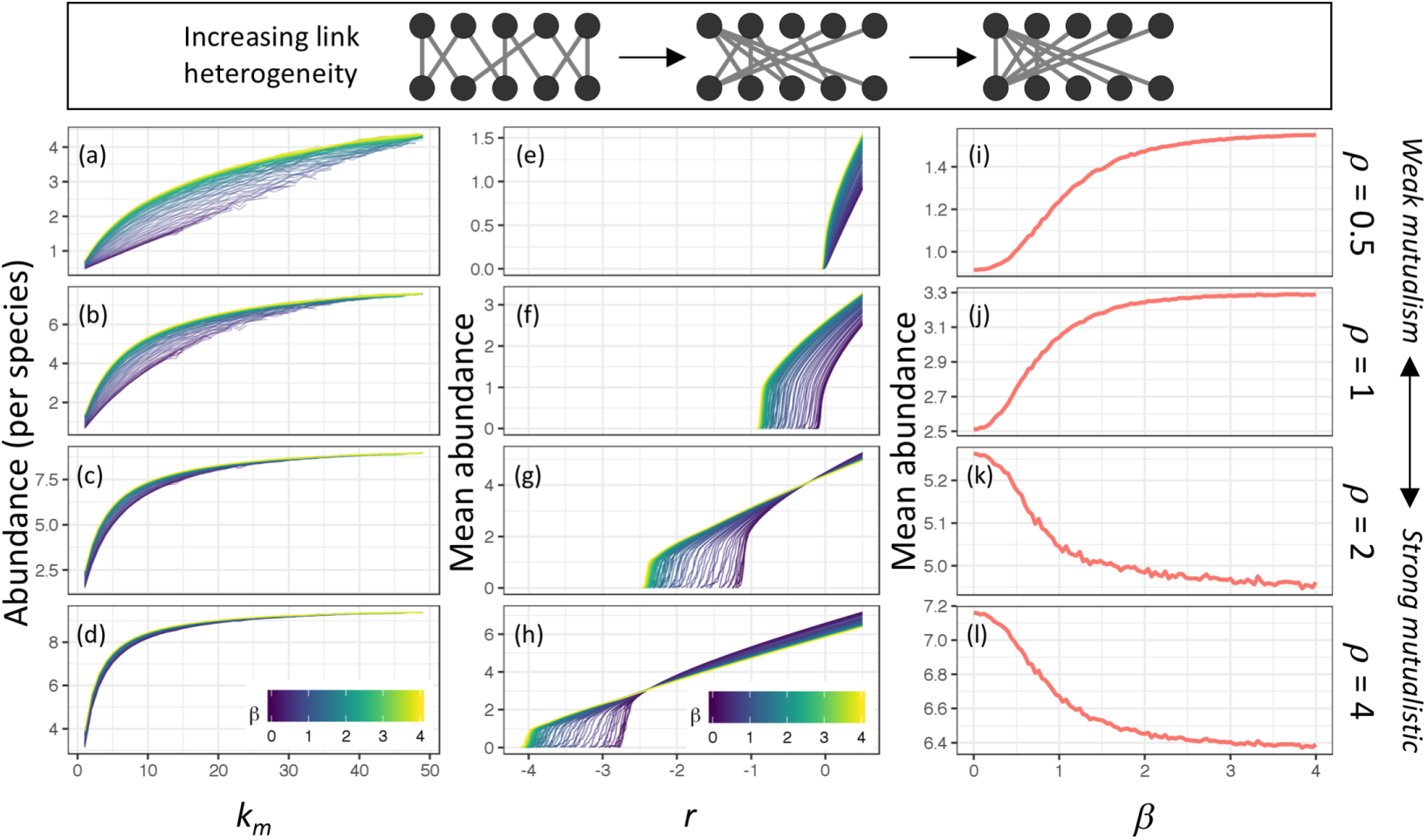
Effect of degree heterogeneity on total abundance. The upper panel shows schematic bipartite mutualistic networks with increasing degree heterogeneity (*β*), as an interpretive aid. (a-d) Abundance of individual species with different numbers of interactions (*k_m_*), shown for different levels of heterogeneity of their interaction network (*β*, colorbar) and relative mutualistic strength *ρ*. (e-h) Dependence of mean abundance on *r*, for different values of heterogeneity (*β*, colorbar) and *ρ*. (i-1) Dependence of mean abundance on heterogeneity (*β*), for each of the different values of *ρ*. Simulation conditions are: *n* = 100, *k_c_* = *n*/2 — 1, *s* = 1, *α* = 3, *h* = 0.1 and the average number of mutualistic interactions 〈*k_m_*〉 = 6, with *ρ* = 0.5,1,2,4 respectively, representing weak to strong mutualism, *r* = 0.5 for (a-d), (i-1). All abundances are the mean of 20 simulations, with 95% quantile of relative standard deviation approximately 0.01, 0.08, 0.015 for (a-d), (e-h), (i-1) respectively.

We find increased degree heterogeneity (larger *β*) has the potential to increase the deviation amongst species abundance, and where this happens there is a reduction in the abundance of the rarest species (ARS). This matters because ARS is strongly negatively correlated with the largest eigenvalue of the relevant Jacobian matrix (**J**), hence determining both the apparent resilience [11] and the occurrence of the first species extinction (which we distinguish from the full collapse of all species). The degree to which heterogeneity drives resilience is however strongly determined by the strength of mutualistic relationships. If mutualism is weak (*ρ* < 1), degree heterogeneity has almost no effect on ARS (or the first extinction (Fig.5a, where *ρ* = 0.5). Under strong mutualism (*ρ* ≥ 1), *r* is found to be a key control of the dependence of ARS on *β* : when *r* is relatively large (sufficient to yield stable finite abundance for all species over the simulated range 0 < *β* < 4), increasing heterogeneity first decreases ARS to a minimum at *β* ≈ 1 then subsequently increases it asymptotically (this response being more exaggerated under stronger mutualism); with progressively lower *r*, the *β*—dependence of ARS becomes more muted, and increased heterogeneity provides no more than a delaying factor for the inevitable loss of species, if *ρ* is reduced to 1. However, in addition to these reductions in apparent resilience, which are essentially determined by changes in **J**, we find a subsequent converse effect which enhances resilience. As degree heterogeneity is increased, a reduction occurs in the largest eigenvalue of the shadow Jacobian matrix **J**˜ (with qualitatively similar form to that of changes in ARS with *β* and *r*, SOM). This serves to increase the resilience of the species that remain following the first extinction, and the persistence of the community under lower values of *r* (e.g. harsher environmental conditions; SOM). This effect increases asymptotically with *β*, and more dramatically under stronger mutualism (larger *ρ*). In light of these findings, a question emerges naturally as to whether trade-offs between abundance and resilience exist in real ecological systems. We see that stronger mutualism (larger *ρ*) provides a benefit in terms of abundance, and under increased heterogeneity (*β* >∼ 2; γ <∼ 1.5) we see greater resilience for both the first and last species extinctions under reduced *r* (harsher conditions). However, higher heterogeneity incurs a ‘penalty’ in terms of abundance under relatively high intrinsic growth rates, and this happens at increasingly lower *r* as the strength of mutualism is increased. For real systems subject to variation in *r*, the necessity for resilience perhaps sets a lower feasible limit for *β* (upper limit of γ ≈ 1.5). Empirical evidence ( [46, 47, 48]) for approx. 1 ≤ γ ≤ 1.25 fits with the notion of parameters which avoid the loss of resilience under higher gamma values, while perhaps hedging against relative losses of abundance due to volatility in environmental conditions (driving variation in *r*). Further work is required to test this proposition. We have explored additional related deviations from the mean field model relating to the trade off between the strength and number of mutualistic interactions, variance in the basic parameters and differences in species numbers of the two interacting groups, details of which are provided in the SOM.

**Figure 5.**
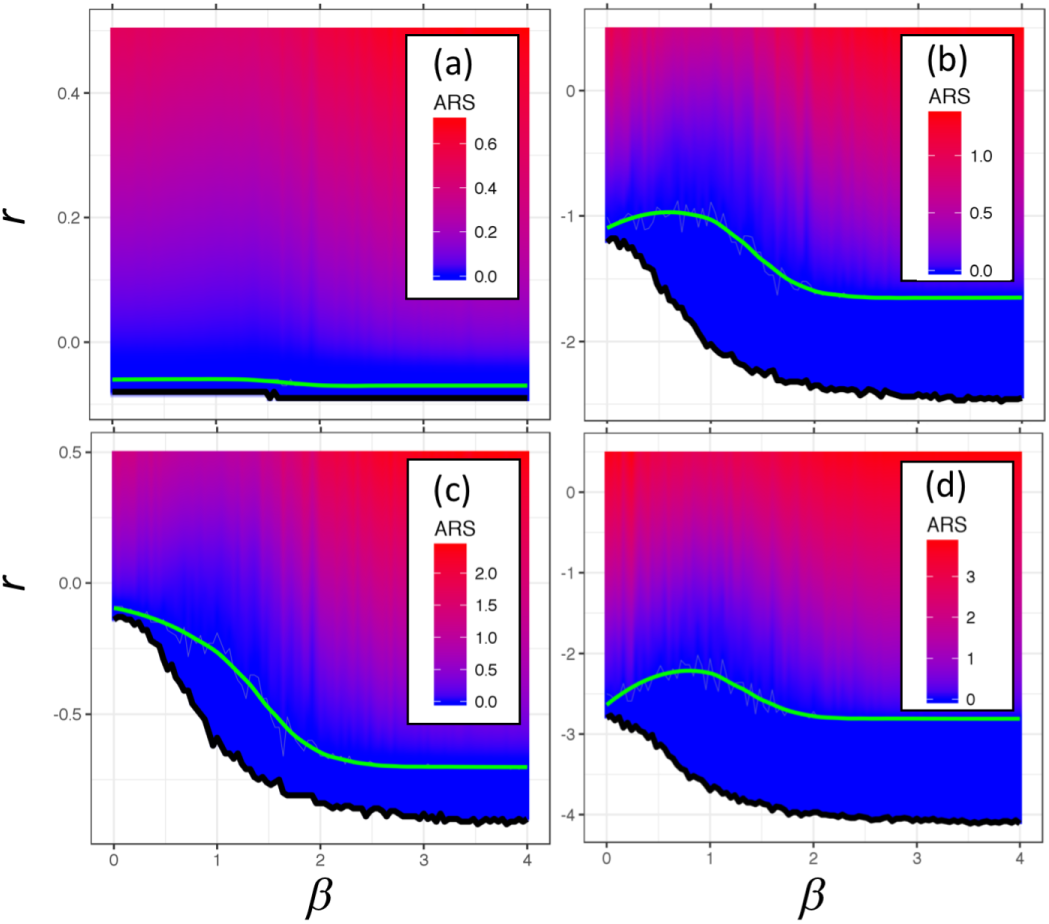
Effect of degree heterogeneity on resilience. Resilience is represented by abundance of the rarest species (ARS). Relationship of ARS (colorbar), first extinction (smoothed green line), and full collapse (black line) to heterogeneity (*β*) under decreasing *r* (Y axis), for (a) *ρ* = 0.5, (b) *ρ* = 1, (c) *ρ* = 2, (d) *ρ* = 4. Simulaton conditions: *n* = 100, *k_c_* = *n*/2 — 1, *s* = 1, *α* α 3, *h* α 0.1 and the average number of mutualistic interactions 〈*k_m_*〉=6. All the ARS values are the mean of 20 simulations (90% quantile of RSDs is around 0.21, 0.83, 0.73, 0.51 respectively).

Deep insights come from connecting basic parameters to system-level phenomena, and in the case of mutualistic systems we see the crucial importance of both non-linear handling-time and mutualistic strength in determining the presence of alternative stable states and the nature of transitions between them. Further, there is potential practical relevance in understanding different types of critical transitions where they occur; in the present context, *splitting transitions* potentially pose greater risks compared to *consistent* transitions, being triggered under less harsh environmental conditions and preceded by hard-to-detect early-warning indicators. The structure of mutualistic interaction networks is seen to play a key role in mediating potential trade-offs between abundance and stability, and this also has potential practical relevance for the conservation of ecosystems and their function. Similar investigations of food-webs and mixed (mutualistic, competitive, exploitative) interaction systems are necessary in order to develop a broad understanding of ecological dynamics, and these are possible using the present theoretical framework.

## METHODS

The General Model We start with the general case of a community composed of n species, divided into two groups: *n_p_* primary producers and *n_a_* species of what can be regarded as animals, such as insects, seed dispersers (*n* = *n_p_* + *n_a_*). Species belonging to the same group are in direct competition, while mutualistic interactions occur between species belonging to the different groups. The deterministic dynamics of the *n* species are described by a system of *n* differential equations, written in matrix form as: 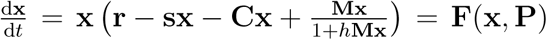, where x is the vector of species abundances, **P** = [**r**, s, **C**, **M**,*h*] is the set of parameters, including vectors of per capita intrinsic growth rates **r** (which can be adjusted to simulate changes in environmental stress), per capita self-regulation strength s, competitive interactions matrix **C**, mutualistic interactions matrix **M**, and handling time *h*. The competitive interaction matrix **C** represents the competitive effects among species within the same group, constructed from the competitive adjacency matrix **G***_c_*. **M** represents the mutualistic effects between species of the different groups, constructed from the mutualistic adjacency matrix **G***_m_*.

We suppose that this mutualistic system has a feasible equilibrium where abundances of all species are strictly positive, hence: 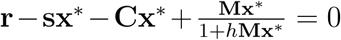, where 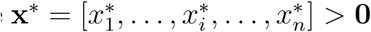 is the vector of equilibrium abundances of all *n* species. The Jacobian matrix at equilibrium can then be written: **J** = diag(x*) · **J**^∼^ = diag(x*) · (—diag(s) — **C** + diag(*ϕ*) · **M**), where *ϕ* = [*ϕ*_1_, · · ·, *ϕ*_*n*_] and 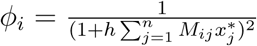 is the ‘effective mutualistic strength’ on species *i* at equilibrium, caused by positive handling time (*h* > 0).

The stochastic dynamics of the n species can be written in matrix form as: dx = **F**(x, **P**)dt + Σ · d**W**, where **F**(x, **P**) represents the deterministic dynamics of each species; d**W** represents a vector of derivatives of independent Wiener processes (Gaussian noise with mean 0 and variance 1); **Σ** is a diagonal matrix with identical diagonal elements equal to σ, representing the variances and covariances of n independent Gaussian noise elements. The variance-covariance matrix **V** of its stationary probability distribution can be approximately obtained by solving the continuous Lyapunov equation equation [49, 50, 51]: **VJ** + **J**^T^**V** = —**Σ**^2^ = —diag(*σ*^2^). This connects the deterministic Jacobian matrix **J** (at equilibrium) with the variance-covariance matrix **V** of the stationary probability distribution for stochastic dynamics.

The Mean Field Model We exclude structural features of the interactions such as degree heterogeneity and modularity, thus all *n* species have an equal number of mutualistic interactions (*k_m_*) and competitive interactions (*k_c_*); variance is excluded from all parameters **P** = [**r**, s, **C**, **M**] using a mean field approximation[8, 32, 28]; and each group contains an equal number of species 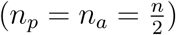 (see SOM for full implications).

Under these conditions the equilibrium condition equation is degenerated to a scalar equation 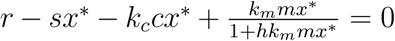, where *x*\* > 0 is the same equilibrium abundance for all *n* species. Transforming this to a quadratic provides two possible equilibrium abundance solutions, where the ‘lower solution’ is always unstable: 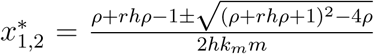, where 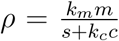 is the ratio of total mutualistic to total competitive strength. The Jacobian matrix at the feasible equilibrium is reduced to: **J** = *x*\*(—*s* — *c***G***_c_* + *ϕ_m_***G**_*m*_), where 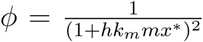 is the reduced version of the ‘effective mutualistic strength’.

The eigenvalue distributions of the Jacobian matrix **J** essentially conform to Allesina and Tang’s theory[6, 7, 52], that the eigenvalues include two parts: a single eigenvalue equal to the (expected) row sum of the matrix, and secondly the bulk of other eigenvalues which are approximately distributed as an semicircle centred at — *E* (the negative expectation of all off-diagonal elements). The eigenvalues of **J** can be calculated from the eigenvalues of the mutualistic and competitive adjacency matraces **G**_*m*_ and **G**_*c*_. (see also the SOM for discussion of divergencies from [6, 7, 52]). Thus, we can define three components of the eigenvalues of the Jacobian matrix. First, the ‘dot’ eigenvalue equal to the row sum, calculated as λ*_d_*(**J**) = *f* (*r, h,ρ*) (see SOM). Second, we name the right most (largest) value in the semicircle as the ‘semicircle’ eigenvalue, defined by: λ*_s_*(**J**) = *x*\*(—*s* + *c* + *ϕm*λ_*S*_(**G**_*m*_)), where λ*_S_*(**G**_*m*_) is the estimated ‘semicircle’ eigenvalue of the mutualistic adjacency matrix (conditioned by n and *k_m_*). Third, we define the difference between the ‘dot’ eigenvalue and the ‘semicircle’ eigenvalue as the ‘spectral gap’ of the Jacobian matrix: Δ∼ = λ*_d_*(**J**) — λ*_S_*(**J**) = *x*_1_*c*(*k_c_* + 1)(—1 + *ϕ*Δ)), where 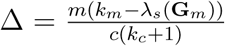is the ratio of the spectral gaps of the mutualistic and competitive interaction matrices. If the spectral gap Δ∼ > 0, i.e. the ‘dot’ eigenvalue is larger than the ‘semicircle’ eigenvalue (therefore also larger than all the eigenvalues in the semicircle), we refer to the ‘dot’ eigenvalue dominating the eigenvalue distribution of the Jacobian matrix. If the spectral gap Δ∼ < 0, the ‘dot’ eigenvalue is ‘submerged’ into the semicircle, and the ‘semicircle’ eigenvalue dominates. Therefore, the largest eigenvalue of **J** is defined λ_1_ (**J**) = max(λ_d_(**J**), λ_*s*_(**J**)). The Jacobian matrix for the mean field model is symmetric, i. e. **J** = **J**^T^ and therefore the variance-covariance matrix **V** with stochastic dynamics simplifies to 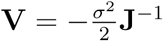.

## References

[1] Ives, A. R. & Carpenter, S. R. Stability and diversity of ecosystems. science 317, 58–62 (2007).

[2] Allesina, S. & Tang, S. The stabilitycomplexity relationship at age 40: a random matrix perspective. Population Ecology 57, 63–75 (2015).

[3] McCann, K. S. The diversity-stability debate. Nature 405, 228–233 (2000).

[4] Battiston, S. et al. Complexity theory and financial regulation. Science 351, 818–819 (2016).

[5] Helbing, D. Globally networked risks and how to respond. Nature 497, 51–59 (2013).

[6] Allesina, S. & Tang, S. Stability criteria for complex ecosystems. Nature 483, 205–208 (2012).

[7] Tang, S., Pawar, S. & Allesina, S. Correlation between interaction strengths drives stability in large ecological networks. Ecology Letters 17, 1094–1100 (2014).

[8] Bastolla, U. et al. The architecture of mutualistic networks minimizes competition and increases biodiversity. Nature 458, 1018–1020 (2009).

[9] James, A., Pitchford, J. W. & Plank, M. J. Disentangling nestedness from models of ecological complexity. Nature 487, 227–230 (2012).

[10] Thbault, E. & Fontaine, C. Stability of ecological communities and the architecture of mutualistic and trophic networks. Science 329, 853–856 (2010).

[11] Suweis, S., Simini, F., Banavar, J. R. & Maritan, A. Emergence of structural and dynamical properties of ecological mutualistic networks. Nature 500, 449–452 (2013).

[12] Beisner, B., Haydon, D. & Cuddington, K. Alternative stable states in ecology. Frontiers in Ecology and the Environment 1, 376–382 (2003).

[13] Kfi, S., Holmgren, M. & Scheffer, M. When can positive interactions cause alternative stable states in ecosystems? Functional Ecology 30, 88–97 (2016).

[14] Scheffer, M., Carpenter, S., Foley, J. A., Folke, C. & Walker, B. Catastrophic shifts in ecosystems. Nature 413, 591–596 (2001).

[15] Scheffer, M. & Carpenter, S. R. Catastrophic regime shifts in ecosystems: linking theory to observation. Trends in Ecology & Evolution 18, 648–656 (2003).

[16] Ives, A. R., Gross, K. & Klug, J. L. Stability and variability in competitive communities. Science 286, 542–544 (1999).

[17] Lehman, C. L. & Tilman, D. Biodiversity, stability, and productivity in competitive communities. The American Naturalist 156, 534–552 (2000).

[18] Thibaut, L. M. & Connolly, S. R. Understanding diversity-stability relationships: towards a unified model of portfolio effects. Ecology Letters 16, 140–150 (2013).

[19] Loreau, M. & Mazancourt, C. Species Synchrony and Its Drivers: Neutral and Nonneutral Community Dynamics in Fluctuating Environments. The American Naturalist 172, E48–E66 (2008).

[20] Gross, K. et al. Species Richness and the Temporal Stability of Biomass Production: A New Analysis of Recent Biodiversity Experiments. The American Naturalist 183, 1–12 (2014).

[21] Scheffer, M. et al. Early-warning signals for critical transitions. Nature 461, 53–59 (2009).

[22] Dakos, V. et al. Methods for Detecting Early Warnings of Critical Transitions in Time Series Illustrated Using Simulated Ecological Data. PLoS ONE 7, e41010 (2012).

[23] Carpenter, S. R. et al. Early Warnings of Regime Shifts: A Whole-Ecosystem Experiment. Science 332, 1079–1082 (2011).

[24] Dakos, V. & Bascompte, J. Critical slowing down as early warning for the onset of collapse in mutualistic communities. Proceedings of the National Academy of Sciences 201406326 (2014).

[25] Naeem, S. & Li, S. Biodiversity enhances ecosystem reliability. Nature 390, 507–509 (1997).

[26] Cardinale, B. J. et al. Biodiversity simultaneously enhances the production and stability of community biomass, but the effects are independent. Ecology 94, 1697–1707 (2013).

[27] Okuyama, T. & Holland, J. N. Network structural properties mediate the stability of mutualistic communities. Ecology Letters 11, 208–216 (2008).

[28] Rohr, R. P., Saavedra, S. & Bascompte, J. On the structural stability of mutualistic systems. Science 345, 1253497 (2014).

[29] Bascompte, J., Jordano, P. & Olesen, J. M. Asymmetric coevolutionary networks facilitate biodiversity maintenance. Science 312, 431–433 (2006).

[30] Vzquez, D. P. et al. Species abundance and asymmetric interaction strength in ecological networks. Oikos 116, 1120–1127 (2007).

[31] Rooney, N., McCann, K., Gellner, G. & Moore, J. C. Structural asymmetry and the stability of diverse food webs. Nature 442, 265–269 (2006).

[32] Saavedra, S., Rohr, R. P., Dakos, V. & Bascompte, J. Estimating the tolerance of species to the effects of global environmental change. Nature Communications 4 (2013).

[33] Lever, J. J., van Nes, E. H., Scheffer, M. & Bascompte, J. The sudden collapse of pollinator communities. Ecology Letters 17, 350–359 (2014).

[34] Grilli, J., Rogers, T. & Allesina, S. Modularity and stability in ecological communities. Nature Communications 7, 12031 (2016).

[35] Ives, A. R. & Hughes, J. B. General relationships between species diversity and stability in competitive systems. The American Naturalist 159, 388–395 (2002).

[36] Otto, S. B., Rall, B. C. & Brose, U. Allometric degree distributions facilitate food-web stability. Nature 450, 1226–1229 (2007).

[37] Allesina, S. et al. Predicting the stability of large structured food webs. Nature Communications 6 (2015).

[38] Mougi, A. & Kondoh, M. Diversity of Interaction Types and Ecological Community Stability. Science 337, 349–351 (2012).

[39] Scheffer, M. et al. Anticipating critical transitions. Science 338, 344–348 (2012).

[40] Gao, J., Barzel, B. & Barabsi, A.-L. Universal resilience patterns in complex networks. Nature 530, 307–312 (2016).

[41] Kuznetsov, Y. A. Elements of applied bifurcation theory, vol. 112 (Springer Science & Business Media, 2013).

[42] Zeeman, E. C. Catastrophe theory (Springer, 1979).

[43] Cubitt, T. S., Perez-Garcia, D. & Wolf, M. M. Undecidability of the spectral gap. Nature 528, 207–211 (2015).

[44] Jonhson, S., Domnguez-Garca, V. & Muoz, M. A. Factors Determining Nestedness in Complex Networks. PLoS ONE 8, e74025 (2013).

[45] Goh, K.-I., Kahng, B. & Kim, D. Universal behavior of load distribution in scale-free networks. Physical Review Letters 87, 278701 (2001).

[46] Montoya, J. M., Pimm, S. L. & Sol, R. V. Ecological networks and their fragility. Nature 442, 259–264 (2006).

[47] Jordano, P., Bascompte, J. & Olesen, J. M. The ecological consequences of complex topology and nested structure in pollination webs. Plant-pollinator interactions: from specialization to generalization 173–199 (2006).

[48] Jordano, P., Bascompte, J. & Olesen, J. M. Invariant properties in coevolutionary networks of plantanimal interactions. Ecology letters 6, 69–81 (2003).

[49] Gardiner, C. W. Handbook of stochastic methods (Springer-Verlag, Berlin, 1985).

[50] Neumaier, A. & Schneider, T. Multivariate autoregressive and Ornstein-Uhlenbeck processes: estimates for order, parameters, spectral information, and confidence regions. ACM Transactions in Mathematical Software (1998).

[51] Suweis, S. & D’Odorico, P. Early Warning Signs in Social-Ecological Networks. PLoS ONE 9, e101851 (2014).

[52] Tang, S. & Allesina, S. Reactivity and stability of large ecosystems. Population Dynamics 2, 21 (2014).

